# Software tools for automated transmission electron dmicroscopy

**DOI:** 10.1101/389502

**Authors:** Martin Schorb, Isabella Haberbosch, Wim JH Hagen, Yannick Schwab, David N Mastronarde

## Abstract

In the recent years, electron microscopy in the life sciences has witnessed increasing demand for high-throughput data collection in both structural and cellular biology. We present a combination of software tools that enable automated acquisition guided by image analysis for a wide variety of Transmission Electron Microscopy applications. Using these tools, we demonstrate dose-reduction in single particle cryo-EM experiments, fully automated acquisition of every single cell in a plastic section and automated targeting of features on serial sections for 3D volume imaging even across multiple grids.

## Motivation/Introduction

As quantitative observations become standard in biological research, there is a growing need to increase the throughput in data acquisition. New technologies such as improved detectors, microscope hard- and software and the increase in compute performance have led to a significant speed-up in Transmission Electron Microscopy (TEM) imaging while at the same time improving data quality^1–3^. The major bottleneck during the acquisition procedure remains the selection of features of interest, namely identifying their exact coordinates for acquisition. This selection procedure is currently done mostly manually. Advanced tools for image analysis exist and already allow for automatic detection of specific features. The approach of employing feedback from image analysis to define the subsequent acquisition is applied in high-throughput fluorescence microscopy^4,5^. In contrast, there is so far no solution to incorporate such techniques into TEM data acquisition workflows.

We have developed two software tools that make automation and feedback approaches available for TEM:

1. *SerialEM*^6^ offers control of the microscope and its imaging detectors in a very flexible manner. In the last years of developing this software, new features for automation have been introduced, that we like to describe in this article. The *Navigator* functionality is the key element to image multiple regions of a specimen with pre-defined acquisition parameters in an automated manner. By using the built-in scripting functionality, users can create and perform highly customized acquisition routines.
2. To provide SerialEM with the necessary positional information for acquisition, we have developed *emTools*, a Python module that integrates common image analysis applications. By running a specimen-specific processing pipeline, we can provide coordinate and image information for automated acquisition using the Navigator. This approach moves the sample-specific manual identification of features of interest to become an image analysis task. Its modularity allows integration into frameworks such as KNIME^7^ where users can incorporate multiple other software tools.

A schematic illustration of the communication, control and data exchange using SerialEM and the emTools module is depicted in Figure 1.

**Figure 1:**
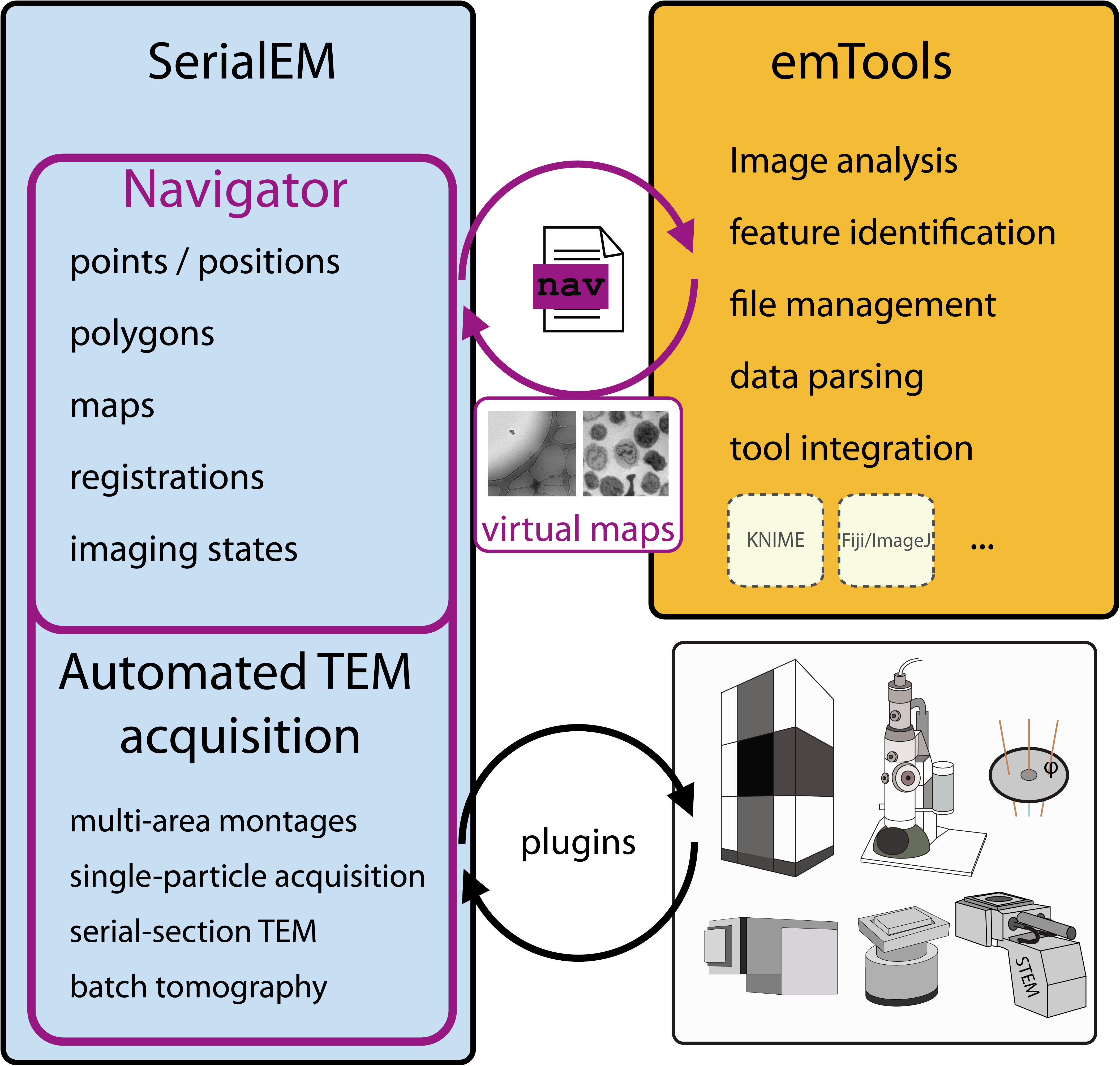
Schematic overview of communication and data transfer between the different components of an automated TEM acquisition. The emTools module serves as a communication interface combining external image analysis tools with SerialEM via the Navigator files and virtual map images.

## Software

### SerialEM

SerialEM^6^ is a versatile software for TEM data collection originally intended mainly for electron tomography acquisition. While its core is made available as open source, it can communicate with microscope and camera hardware of various manufacturers using a plugin scheme. Data presented in this article was acquired using the following microscopes: JEM 2100Plus (JEOL Ltd., Akishima, Japan), Tecnai F30 and Titan Krios G3 (Thermo Fischer Scientific, Waltham, MA, USA). Over the years, the software has received significant additional functionality to increase automation. SerialEM comes with a built-in scripting feature. The users can control and modify most of its functionality using scripts and thus highly individualize their acquisition procedures. In this paragraph, we will explain the main components of SerialEM required for automated acquisition. These functions have been developed over the last few years since the intital description of the software^6^, but have not been described in a scientific publication thereafter.

#### Navigator

The *Navigator* module is the main tool for finding, keeping track of, and positioning at targets of interest. The stage positions of single points can be recorded either by storing stage coordinates while browsing the specimen, or by marking points on an image. The Navigator also stores information for two other kinds of items: *maps* and *polygons*.

#### Maps

A *map* is a specific image for which the Navigator remembers its storage location, the corresponding acquisition parameters for microscope and camera, and how to display the image to the user. Maps are generally used to provide an overview for picking acquisition targets, but also play an important role in allowing repositioning at a target during batch acquisition.

#### Polygons

*Polygons* are connected points drawn on an image to outline an area. Their main use is to define the area to be acquired with a *montage* at a higher magnification. The program can then define optimal parameters and acquire a tiled montage that will cover the requested area.

#### Montages

In case the field of view of a single image is not sufficient to cover the area of interest, SerialEM can acquire a *Montage* of tiles that overlap at their edges. Montages are always taken at regular spacing in a rectangular pattern, but acquisition at tiles which are not needed to fill a polygon can be skipped (Fig. 4A). For larger areas where it is inconvenient to draw a polygon, such as a ribbon of sections, one can move the stage to each corner of the area and record a *corner point* there. The program can then fit a montage to that set of corners points just as for a polygon.

SerialEM’s ability to acquire a montage of overlapping images has been optimized for several different situations. For tilt series acquisition, the different positions are reached with electronic image shift. Larger areas can be acquired with stage movement instead; this method is typically used for acquisitions from large areas, such as serial sections. The program has a set of features, such as periodic focusing, to support acquisition of high-quality images from arbitrarily large areas^8^. Another common use of montages is to acquire overview images for finding targets for data collection; to support this use for cryo-EM, montaging can be done with two of the lower magnification states defined as part of the program’s *low dose* mode.

#### Realign to Item

To reposition the acquisition area accurately at a marked position, the program has a procedure, *Realign to Item*, that takes images under the same conditions as those used to acquire a map and cross-correlates those images with the map. The map needs to be sufficiently larger than the errors in stage positioning that can accumulate over the course of data acquisition; a field of view of at least 10 µm is recommended. A map used for realigning can be either a single image covering sufficient area, or a montage in which the individual tiles need not be so large. After aligning to this map, the realignment routine can also align to a second map acquired at higher magnification; a pair of maps is frequently used to reposition accurately at a defined target area for tilt series acquisition. Using a montage for realignment is efficient when an overview montage is used for initially finding targets, but for other situations the program provides a convenient alternative, the *Anchor Map*. After taking a pair of maps at the first target, the user selects an option to indicate that the lower-magnification map is the template for how to take anchor maps. Thereafter, the user can take an image of the target area at the acquisition magnification and press the“Anchor Map”button; the program will save that image as a map, will then go to the lower magnification and take and save the anchor map. Aside from convenience, this method handles any misalignment between the magnifications by incorporating the misalignment into a displacement between the high magnification map and the anchor map, where the feature of interest does not need to be centered. Overcoming such misalignments is more difficult with a montage map, unless it is taken in SerialEM’s *low dose* mode, where the user can bring images taken at the lower magnification (*View* mode) into alignment with those at the higher one (*Record* mode) with a dedicated image shift control panel.

When using SerialEM to acquire images for single-particle reconstruction, it is routine to acquire montages of promising grid squares in View mode. The next task is to add Navigator points in the centers of selected holes in the carbon support film. Although SerialEM lacks a tool for automatically selecting holes, it does allow one to mark five points to define both the extent and spacing of a set of holes, and then to add a regular grid of points in the adjacent holes. Additional points can then be added, or unwanted points selected and removed easily with the mouse. Larger specimen regions e.g. areas of crystalline or thick ice can be excluded by masking them with polygons.

If a template image of a single ice-covered hole of the support film is provided, this can be used for better centering each hole^9^.

#### Batch acquisition

Batch acquisition by the Navigator is initiated with the *Acquire at Points* dialog, which can acquire an image, a map, or a tilt series or alternatively run a script at each item marked for acquisition. Single-particle acquisition is generally run exclusively through scripts. The dialog offers a choice of operations to be run before acquisition, such as focusing or centering the beam. The *Realign to Item* routine will typically be chosen to guarantee accurate repositioning.

#### Registrations

The Navigator stores a *Registration* number for each item, where items with the same value are considered to belong to the same coordinate system. When a grid is rotated, or removed and reinserted, the coordinate system has changed, and any new Navigator items need to be assigned a different registration number, whereas existing ones need to have their coordinates transformed into the new registration. Similarly, positions imported from a light microscope are assigned a different registration number. To transform coordinates from one registration to another, the user marks a set of points that correspond between the registrations (e.g., features that can be identified in both an imported Light Microscopy image and an EM map). The program can then solve for a shift, rotation, or a linear transformation depending on the number of landmark points and transform coordinates into the current registration. Alternatively, one can save an image before rotating the grid for dual-axis tomography, take a corresponding image afterwards, and the program can find the rotation and shift that aligns the images by cross-correlation. This procedure is called *Align with Rotation*. The resulting transformation is accurate enough over a more limited range of ∼500 µm. Transforming a registration also leads to SerialEM transforming the map images accordingly before using them for *Realign to Item*. This capability allows a transfer of an entire experiment from one transmission electron microscope to another. A single transformation using landmark features then enables one to use not only the previously recorded coordinates but also the maps for subsequent acquisition with an entirely different microscope.

#### “Virtual” map templates

As long as the necessary operating condition for the electron microscope are available as a map item, a template image file with corresponding map parameters can be manually added to a Navigator file. SerialEM is then able to use this information to align to similar features on the current specimen. This procedure can be used to precisely realign to a specific cell in neighboring serial sections. We therefore provide a map an image of that cell on the initial section and use it to identify and target the same cell on the following section (Application 3, Fig. 4C).

One key concept of our automation approach is to provide SerialEM with a *virtual* map that is defined for each acquisition item. With the information about the visual features of the target object in the associated image, an accurately positioned acquisition can be guaranteed through the *Realign To Item* procedure. We acquire a single map with the desired acquisition settings anywhere on the specimen. Its Navigator entry will then be used as a template containing all important parameters other than position and the associated image.

The software SerialEM and all documentation are available at http://bio3d.colorado.edu/SerialEM/

### The emTools module

The module is an open-source collection of Python functions that can interpret and generate SerialEM’s Navigator files, modify Navigator items and perform or trigger image analysis tasks on maps. It is intended to interface between SerialEM – hence the microscope – and any software tool that can run the image analysis and processing necessary for a specific experiment. We have put a strong focus on making this interface as modular as possible to leave maximum freedom in the choice of external tools. The processing can be done at the microscope for immediate feedback to the acquisition or offline for complex tasks that require powerful computing resources.

A general procedure when using emTools would be to first read and parse a Navigator file and identify the image file(s) of the desired map(s). SerialEM first tries to locate maps based on the absolute storage location of the associated image available in the Navigator file. In case the file is not found, it will try relative paths. Therefore it is usually possible to simply transfer the entire data directory including the Navigator file and all associated image data to a central storage and perform any processing on powerful computers. The interpretation of the relative paths allows to then re-insert all data back into SerialEM. In order to mark the maps for processing, we use the “*Acquire*” flag in the Navigator.

In order to process a map consisting of multiple tiles (i.e. a *Montage*) the emTools module’s function *mergemap* will make use of stitching routines available in the IMOD^10^ software package. It then imports the resulting single stitched image and makes it available for image processing. It also takes care of preserving the Navigator’s coordinate system for that map, so any items identified or created by other scripts can be placed correctly.

Once done with the processing and generating or modifying all affected Navigator items the emTools functions fill an output Navigator file that is then used as a basis for further acquisition.

We have applied emTools workflows to automate TEM acquisition either running as standalone Python scripts or embedded into a KNIME workflow. This generic data analysis software^7^, in addition to built-in capabilities^11^, can easily incorporate image processing modules from various sources and different programming languages and run them as part of a workflow. We use this approach because some image processing functionalities are readily available using special software tools such as ImageJ^12,13^, CellProfiler^14^, R^15^, Ilastik^16^ or MATLAB compared to limited implementations in Python. The modular approach of the emTools functions combined with the KNIME framework allow incorporating virtually any type of already existing image processing routine to identify features of interest.

The main functions of the emTools module comprise tools to:

- parse files in the SerialEM/IMOD “autodoc” format, in particular Navigator files
- extract coordinate and image information from Navigator files and modify these
- handle Navigator items as Python dictionaries
- merge mosaic maps (tiles) using IMOD to obtain smooth merged images
- enable any general image processing pipeline to identify features of interest
- generate Navigator files with coordinates determined by an image analysis pipeline
- generate maps with pre-defined microscope settings from a map template and incorporate external image data
- filter Navigator files to only keep a selection of items
- duplicate selected Navigator items into a new registration

The emTools module uses functionality of the following Python packages: tifffile, numpy^17^, scipy^17^, scikit-image^18^, mrcfile^19,20^, and pandas^21^ for KNIME integration.

The emTools module is available for download or further collaborative development at https://git.embl.de/schorb/serialem_tools.

## Applications

### 1. High yield automated cryo-EM data acquisition of complicated samples

To enable averaging of images and thus achieve high-resolution structural information, single particle TEM acquisition aims at acquiring many (multiple thousand) high magnification images of thin ice areas containing particles of interest. Typically, either holey carbon grids with a periodic pattern of holes or grids coated with lacey carbon are used.

During an automated single particle acquisition using SerialEM, we usually employ the following procedure:

#### Grid map

We acquire a low magnification (e.g. 100×, 6×6) montage that covers the entire holey carbon TEM grid. In this montage, grid squares of interest are identified (Fig. 2A).

**Figure 2:**
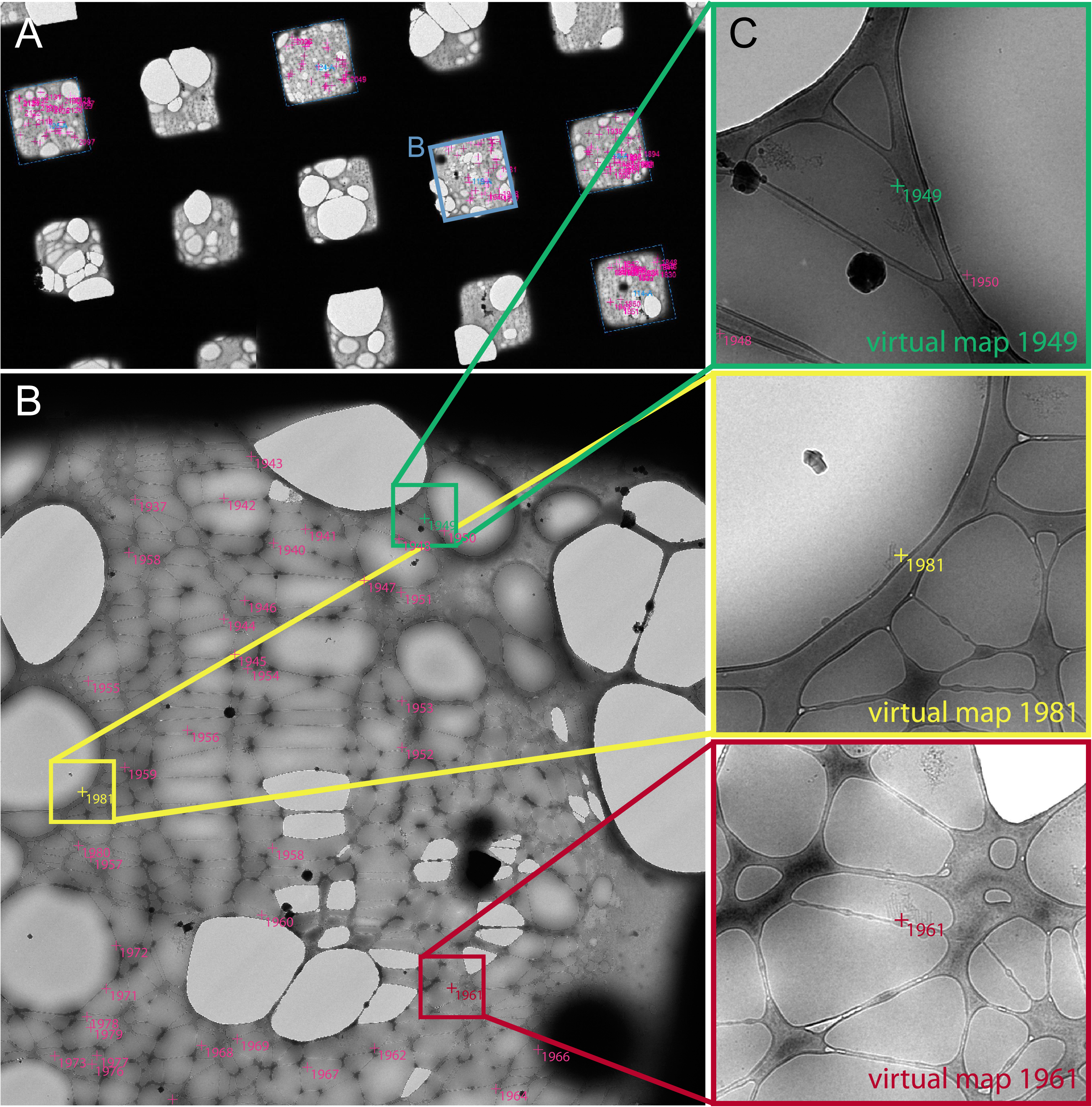
Example application of the automatic generation of virtual maps to acquire TORC1 filamentous particles^23^ with cryo-EM. **A** – part of the overview map of the entire EM grid. Blue boxes indicate the location of higher-magnification maps. The mapped grid square shown in B is marked. The magenta crosses mark the points of interest that were manually chosen in the maps of each grid-square. **B** – 2×2 montage map covering a single grid square at 2250× with a pixel size of 6.2 nm without binning. The colour marks indicate the target positions that were manually clicked within this map. Coloured boxes and marks indicate the example regions where the corresponding example virtual maps were extracted. **C** – Three example virtual maps that the script extracted from the map shown in B. They are used to realign at high magnification (15 000×) to the target positions. Scale: blue box in A, FOV of B: 47 µm; coloured boxes in B, FOV of C: 3.8 µm.

#### Grid square maps

For each identified grid square, we acquire a montage (“grid square map”) at medium magnification (e.g. 2000×, 2×2) to provide more visible detail (Fig. 2B.).

#### Selecting acquisition areas

Larger particles (e.g. filaments or viruses) can be identified in these montages, but the user needs to screen at higher magnification which holes contain smaller particles not visible in these montages. For all holes containing evenly distributed large or small particles, Navigator points are added either manually or using SerialEM’s capability to generate an evenly-spaced grid of points.

#### Automated acquisition

Automatic acquisition is run by executing a script on each previously selected Navigator point. The script moves the sample stage to the hole and centers the hole at medium-high magnification (e.g. 15 000×) acquisition by aligning to the template image of a centered hole that the user stores in a dedicated alignment buffer. It then performs optional autofocus and drift measurements to be followed by acquiring multiple images in a regular circular pattern around the center of the hole at high magnification (e.g. 100 000×) using image shift. This script is available at the online repository^22^.

#### Problematic samples

For many samples, particle concentration and spreading can be a problem. Acquiring such samples using a predefined acquisition template for each hole results in a low yield of useable images for further processing. Considering the high costs of running cryo-transmission electron microscopes this is also uneconomic.

When such problematic samples contain large particles on continuous or lacey carbon grids that are visible in the grid square maps at medium magnification, one conventional option is to use SerialEM’s *Realign to Item* before running the high-magnification acquisition template. The large field of view of the medium magnification images however leads to overlap of imaged regions. This increases the total applied exposure dose and thus cause radiation damage and reduces the quality of the final high magnification images.

The second conventional option is to manually select all particles of interest in the medium magnification maps and acquire another round of single images at medium-high magnification (15 000×) in Low-Dose View mode, saved as maps. In these maps, particles are then selected once again and automated acquisition is started using these maps for the *Realign to Item* procedure. This workflow is efficient for both holey and lacey grids but requires additional setup and significant mapping time and also introduces extra dose at each position.

#### Solution for problematic samples or when using lacey grids

After adding Navigator points for particles of interest in the medium magnification grid square maps and marking them for acquisition, the emTools script *maps_aquire.py* is run on the Navigator file. The script uses IMOD to merge the grid square maps linked to the selected Navigator points if necessary. It then extracts virtual medium-high magnification maps at each position with the parameters from a medium-high magnification template map taken with the SerialEM Low-Dose View parameters. It crops the respective image areas from the medium magnification grid square maps for each selected Navigator point and transforms the extracted images. The script accounts for both the relative scaling and rotation between the grid square map and the medium-high magnification template and thus makes the extracted image suitable for direct correlation with an image acquired at medium-high magnification.

As result, the script generates a new Navigator file containing all these virtual maps and marks these maps for acquisition forcing SerialEM to realign to each of the virtual maps.

The cryo-EM workflow then becomes:

- Acquire grid map.
- Acquire grid square maps.
- Select acquisition areas by adding Navigator points in the grid square maps.
- Setup SerialEM Low-Dose, acquire a Low-Dose View image, save it as a map.
- Run emTools script *maps_aquire.py* on the saved Navigator file.
- Load the new Navigator file generated by *maps_aquire.py*.
- Start automated acquisition.

Using emTools for the creation of virtual medium-high magnification maps directly from the grid square maps enables automated high-yield acquisition of cryo-EM data with minimal dose on samples with low concentrations of large particles, using either continuous, holey or lacey carbon grids.

In the presented example workflow, we target TORC1 filaments^23^ for single particle cryo-EM acquisition. Examples of the virtual maps generated by the script are shown in Figure 2C.

We routinely employ the acquisition strategies described above at EMBL’s Cryo-Electron Microscopy Service Platform^24^.

### 2. systematic acquisition of individual cells on resin sections using feedback TEM

A few organelles, such as centrosomes, are present as single copies in a given cell. In a typical TEM analysis, the probability to observe a centriole on a 200 nm section of resin-embedded human leukocytes^25^ ranges between 3 to 5%. In order to study the ultrastructure of centrosome aberrations in these cells, we have developed a workflow to automatically acquire each individual cell on serial plastic sections. With dimensions of about 500×200 µm, these sections would typically display from several hundred up to 1500 cells.

First, we acquire a low-magnification map of a single section (200× - 400×) and select it for acquisition. This will trigger the emTools scripts to process this map. We also acquire a single map at an intermediate magnification that will be used as a template for the virtual maps the scripts will generate for each detected cell.

We then run a specimen- or phenotype-specific image analysis pipeline (Fig. 3B) that segments individual objects of interest in the image, in this case cross-sections through cells, and returns an image file with labels for each object. We use the KNIME platform to link and incorporate various independent tools for image analysis (in our case Python and ImageJ/Fiji). The user can easily control image analysis parameters through a GUI dialog without the need of having knowledge in programming. Figure 3A shows the layout of the KNIME workflow and the graphical controls where users can modify a few selected parameters.

**Figure 3:**
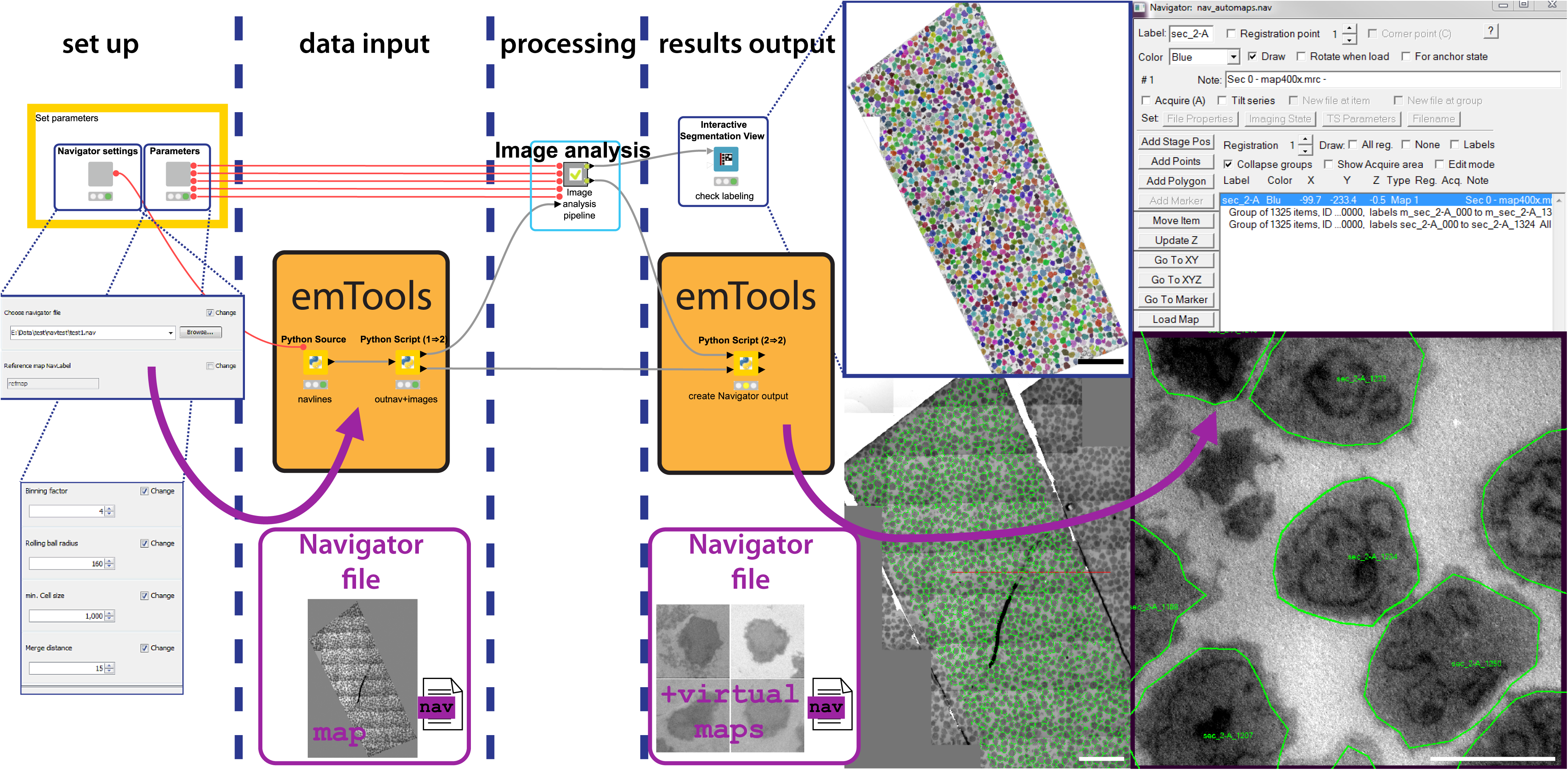
Illustration of the workflow to automatically acquire all cells on a resin section. The left side shows the KNIME workflow with nodes and data connectors. Dark blue boxes show the configuration dialogs where the user provides the location of the Navigator file and some parameters for image analysis. The initial Python nodes use emTools to parse the Navigator file and import the map image(s). The image analysis node contains the entire procedure that extracts the position and outline of each individual cell from the image. Its detailed contents are depicted in Supplementary Fig. S1. The user can monitor the result of the segmentation using the “Interactive Segmentation View” node. The resulting Navigator file that is generated by the final Python scripting node contains a virtual map and the polygon outline for each cell. On the bottom left of the results panel, the low-magnification input map is displayed with all polygons drawn on top. The Navigator window with the item groups of maps and associated polygons is shown at the top-right. An example virtual map is shown below. Scale bars: 50 µm (overviews), 5 µm (virtual map).

In our example, the workflow will employ emTools to:

- merge the map(s) using IMOD (if mosaic)
- load the map image(s) into the KNIME environment
- feed the images into the analysis pipeline that incorporates tasks run by Python, native KNIME nodes and ImageJ/Fiji to identify the single cells
- extract the outline of each cell and generate polygons
- extract virtual maps at each position with the parameters from the provided template map
- link the polygons to the template maps
- generate a Navigator file with all virtual maps and polygons, select and mark those for acquisition

The result of running the procedure is a Navigator file that for each detected cell contains a virtual map and a polygon defining its contour. The resulting overview in SerialEM and some of the virtual maps centered on the respective cells is shown in Figure 3C.

In the resulting Navigator file, all polygons are already marked for acquisition. In order to initiate an automated run and image each cell, the user only has to simply set the desired microscope condition and launch the *Acquire at points* procedure.

The described procedure can be used to acquire an image stack containing one image per cell. This stack can then be screened easily to categorize cells according to cell type, phenotype or by identifying aberrant morphology. Cells can also be selected upon presence of a rare ultrastructural feature. In the presented example, the software automatically detected 1325 cells. By manually screening the image stack we selected the cells that displayed parts of centrioles for further imaging (see below). The entire image stack is presented in supplementary movie 1.

### 3. Automated serial-section TEM

In a serial-section TEM experiment, one wants to follow a feature across a series of sections that lie adjacent on the TEM grid^26^. In our example, we have selected 60 human leukocyte cells on one of the serial sections (thickness 200 nm) as described above. We now want to acquire the exact same cells on all other sections collected as a ribbon on the grid (Fig. 4A) as well as on the following sections on consecutive grids and thus follow the cells in 3D.

**Figure 4:**
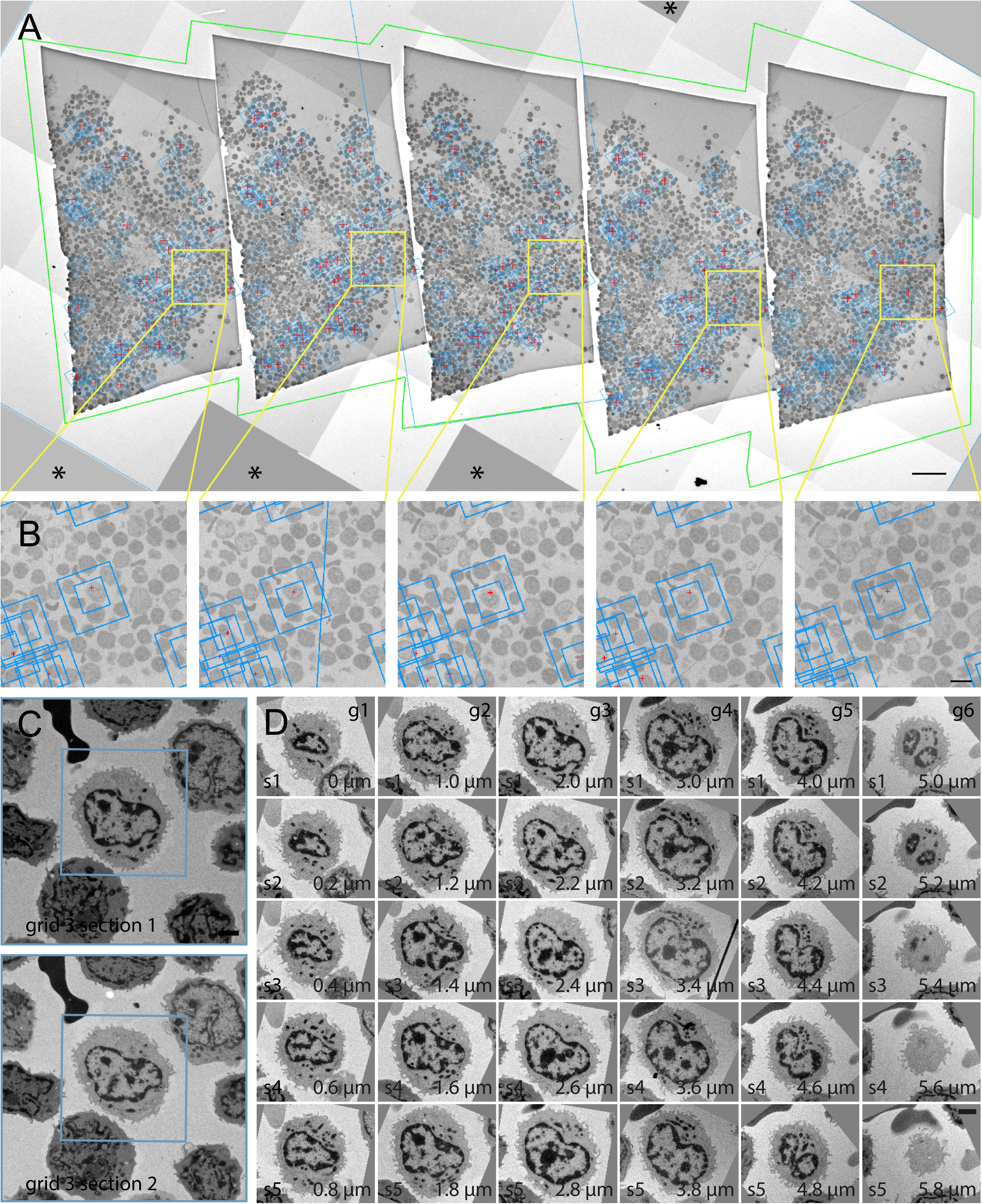
Automated acquisition of cells on serial sections. **A –** Overview of a ribbon of sections placed on a slot grid. The green outline marks the polygon used to acquire this montage map. The blue boxes denote the location of the maps used to realign to each cell. The smaller boxes inside mark higher magnification maps for more precise targeting. Red crosses indicate positions for desired tomography acquisition. **B –** Magnified regions in A showing the location of one cell of interest. **C –** Two maps of a single cell on neighbouring sections. The smaller blue square represents a higher-magnification map to precisely position the final tomogram. **D** – Gallery of images of an individual cell across 30 sections (thickness: 200 nm) spread across 6 grids. The images were automatically aligned using TrakEM2^32^. Another cell that spans cross 9 sections is shown in SM1. Scales: A: 50 µm, B: 20 µm, C/D: 2µm

The procedure we employ is as follows.

For each cell of interest on the initial section, we create a map at a moderate magnification (2300×) with a sufficiently large field of view for realignment. We then select the Navigator items of all interesting cells we like to propagate to the next section, in this case the “target” section, for acquisition.

The emTools script *duplicate_items* then duplicates these map items into a new registration. In order to move them to the target section, we select a few easily recognizable cells on both the initial and on the target section and use these features as *Registration Points*. Taking overview montages at low magnification (∼300×) can help finding cells back and speeds up the process. We then apply the resulting transformation to all duplicated map items. These items are now placed approximately at the correct position on the target section. The few microns of shift can be corrected using the *Realign To Item* procedure when the cellular features provide sufficient visual similarities between adjacent sections.

This is done by running a SerialEM script which then acquires new maps for the target section and labels the Navigator entries such that they match the source Navigator labels. This keeps the indexing of the cells consistent to avoid confusing the many navigator items (cell43_s1, cell43_s2, etc.).

The entire relocation procedure is then repeated for each transition to a neighbouring section. We usually start from a single section where we identify the target cells and then follow them section-by-section in both directions.

In general, objects like cells or tissues whose features do not change dramatically from one section to the following are suitable for this alignment procedure.

In our example, we used this approach to successfully follow each of 60 cells through a series of 45 adjacent serial sections across 9 grids. During a single transition, usually every cell is correctly found and imaged by the described procedure. This applies as well when changing to another grid to image the next set of sections aligning the first section on the new grid with the last on the previous grid. Problems in identifying the correct feature only occur, when the edge of a section, a crack or a fold or large contaminations are present in the field of view. In this case, when usually one or two individual cells are not identified correctly, we manually correct the affected map(s). When the similarity between sections is big enough, such as it is the case for these cells, it can even be sufficient to align all sections on a grid to the template maps from the first section and run the acquisition of new maps for the entire grid at once. For 60 cells, this procedure takes approximately four hours. An example of aligned images of a cell across 42 serial sections spread on 9 grids is shown in Supplementary Movie 2.

In addition to obtaining the full volume of the cells, we were interested in resolving the structure of their respective centrosomes at higher resolutions. For this, we acquired tilt series for each feature of interest can be easily acquired on the consecutive sections using SerialEM batch tomography acquisition. We automatically processed the large amount of acquired tilt series using IMOD’s batch tomogram reconstruction^27^ on a high-performance compute cluster resulting in a full volume of the desired object. An example of such a reconstructed tomographic volume is shown in Supplementary Movie 3.

We have made available the SerialEM scripts used in this experiment at the public Script Repository^28^.

## Discussion

We have developed both software packages described in this article with a strong focus on flexibility. SerialEM is capable of controlling transmission electron microscopes, accessories and detectors from a range of different vendors and covers acquisition strategies like automated single particle acquisition, large 2D mosaic tiling as well as batch tomography on serial sections. The emTools module aims at universally linking SerialEM with a variety of software options to perform image analysis. This means that the main task in setting up an automated TEM workflow for a specific application is now reduced to developing a suitable image analysis routine. These routines are obviously not limited to the examples described in this article, but could also include advanced feature detection using machine learning^16,29^ including convoluted neuronal networks^30,31^. These approaches will also help in reducing the need for manual identification of particles in cryo-EM by automatically picking holes or large particles.

The possibility of imaging every entity of a specimen in a controlled manner offers entirely new possibilities for TEM. Automated classification into different cell types and phenotypes or detection of aberrant morphology can be performed by computational analysis when having large amounts of comparable data at hands. Automation of serial-section TEM significantly reduces the work load required for this technique and opens new possibilities to perform large-scale 3D-EM while maintaining a high lateral resolution. The increased throughput and thus the resulting capability of generating a lot more data that can subsequently be fed into automated analysis pipelines opens possibilities for entirely new types of quantitative TEM experiments.

## Acknowledgements

We thank Ambroise Desfosses and Manoel Prouteau for providing specimens to test and apply the cryo-EM workflow. We acknowledge support from Colin Palmer for mrcfile, Christian Dietz and Clemens von Schwerin for KNIME Image Processing and Python bindings. We like to thank all staff of the EM Core Facility at EMBL for helpful discussions and ideas. We also acknowledge support from the EMBL Center of Bioimage Analysis (CBA). Work on SerialEM was supported originally by grants from NIH and more recently by contributions from users and from JEOL USA, Inc, and by payments for specific projects by Hitachi High Technologies America, Inc, JEOL USA, Inc., and Direct Electron, LP. I.H. is the recipient of a HRCMM (Heidelberg Research Center for Molecular Medicine) Career Development Fellowship.

## Author contributions

M.S., I.H., W.J.H.H and Y.S. designed the experimental applications. M.S. and D.N.M. did the software development. I.H., W.J.H.H. and M.S. collected the data. All authors contributed valuable suggestions to the necessary software functionality and edited the article. M.S. and D.N.M wrote the initial manuscript.

---

**Supplementary Figure S1:** The KNIME image analysis pipeline executed within the meta-node (Fig.3). Each node performs an individual processing task. The respective result is shown above for reference. The output is an image with a distinct pixel intensity value (“label”) for each detected cell.

**Supplementary Movie M1:** The image stack with one overview image for each of the automatically detected 1325 cells. Scale bar: 1 µm.

**Supplementary Movie M2:** Series of Map images of a single cell of interest across 42 serial sections on 9 consecutive grids. The centriole is marked with a blue arrow. Scale bar: 1 µm.

**Supplementary Movie M3:** Reconstruction of a serial tomogram of the centriole region marked in SM1. The acquisition area spans across 6 sections covering approximately 1.2 µm. Scale bar: 200 nm.

